# Situating the Left-Lateralized Language Network in the Broader Organization of Multiple Specialized Large-Scale Distributed Networks

**DOI:** 10.1101/2019.12.11.873174

**Authors:** Rodrigo M. Braga, Lauren M. DiNicola, Randy L. Buckner

## Abstract

Using procedures optimized to explore network organization within the individual, the topography of a candidate language network was characterized and situated within the broader context of adjacent networks. The candidate network was first identified using functional connectivity and replicated across individuals, datasets, acquisition tasks, and analytic methods. In addition to classical language regions near to perisylvian cortex and temporal pole, additional regions were observed in dorsal posterior cingulate, midcingulate, anterior superior frontal and inferior temporal cortex. The candidate network was selectively activated when processing meaningful (as contrast to non-word) sentences, while spatially adjacent networks showed minimal or even decreased activity. Examined in relation to adjacent networks, the topography of the language network was found to parallel the motif of other association networks including the transmodal association networks linked to theory of mind and episodic remembering (often collectively called the default network). The several networks contained juxtaposed regions in multiple association zones. Outside of these juxtaposed higher-order networks, we further noted a distinct frontotemporal network situated between language regions and a frontal orofacial motor region and a temporal auditory region. A possibility is that these functionally-related sensorimotor regions might anchor specialization of neighboring association regions that develop into the language network. What is most striking is that the canonical language network appears to be just one of multiple similarly organized, differentially specialized distributed networks that populate the evolutionarily expanded zones of human association cortex.

## Introduction

The association cortex comprises a mosaic of distributed networks that each interconnect regions in prefrontal, parietal, temporal and midline cortices (Goldman-Rakic 1988; see also Mesulam 1981; 1990). The distributed spatial motif is shared across neighboring regions, leading to a parallel organization of networks (Yeo et al. 2011; Power et al. 2011; Margulies et al. 2016; Braga and Buckner 2017). One hypothesis is that the broad organization of higher-order association cortex is established early in development, with subsequent specialization of cortical zones into distinct networks by activity-dependent processes (Buckner and DiNicola 2019). While much work has previously estimated the spatial relations between multiple higher-order association networks (e.g., Margulies et al. 2016), less emphasis has been placed specifically on the canonical distributed network specialized for language, leaving open the questions of i) whether the language network possesses features similar to other distributed association networks, and ii) how the language network fits within the spatial macroscale organization of the cerebral cortex. Thus, there is a gap in our understanding of how to situate language-responsive regions within the broad mosaic of networks that populate association cortex.

The gap is particularly notable given that the description of interconnected anterior and posterior language regions inspires much of the contemporary study of human brain networks. The classic perisylvian language system, initially identified through case studies of patients with aphasia, included an extended region encompassing inferior frontal gyrus (IFG) just rostral to motor cortex (i.e., Broca’s area) and the posterior superior temporal cortex (pSTC; i.e., Wernicke’s area; Geschwind 1970). Quantitative lesion mapping (Bates et al. 2003; Mirman et al. 2015) and study of progressive aphasia (Mesulam et al. 2013) have highlighted the importance of rostral regions of temporal association cortex extending to the temporal pole (TP), and additional regions at and around traditionally defined Broca’s area. Taken together, classical and contemporary findings on the anatomy of language function support a specialized, left-lateralized network that involves multiple distributed anterior and posterior association regions.

Task-activation studies of language based on group-averages yield an estimate of regions involved in language function that is in many ways consistent with the clinical literature (e.g., Petersen et al. 1988; Wise et al. 1991; Blank et al. 2002; Hickok and Poeppel 2007; Ferstl et al. 2008). Findings converge on a left-lateralized network active during speech reception and production, with regions distributed in anterior and posterior zones that include pSTC often extending rostrally to the temporal pole, and prefrontal regions prominently including the IFG. It is intriguing that this canonical set of language regions broadly adheres to the general motif of other association networks (Goldman-Rakic 1988; Yeo et al. 2011; Power et al. 2011; Margulies et al. 2016). Further, the proximity of language regions in the IFG and temporal association cortex to orofacial motor and auditory cortices (Geschwind 1970; Krubitzer 2007) may have relevance to the development of language pathways (see also Hickok and Poeppel 2007). A core goal of the present work is to use within-individual neuroimaging approaches to characterize the detailed spatial organization of the language network in relation to other nearby functional regions.

A further motivation for exploring the detailed organization of the language network is that group-based studies frequently reveal that the same (or nearby) IFG regions are activated by both linguistic and non-linguistic task demands. This observation led to suggestions that certain parts of the estimated language system act as domain-flexible resources supporting controlled processing (e.g., Thompson-Schill et al. 1997; Poldrack et al. 1999; Gold and Buckner 2002; Burianova and Grady 2007; Hein and Knight 2008; see Blumstein and Amso 2013 for relevant discussion). One possibility is that distinct regions are blurred together in group-averaged data. Of critical importance, when functional zones are defined within individuals, distinct language-specific regions can be defined within IFG that lay in close proximity to, and are surrounded by, less domain-specialized regions (Fedorenko et al. 2010; 2012) that are typically considered part of a separate system called the ‘multiple-demand’ or ‘frontoparietal control network’ (FPN; Duncan et al. 2010; Vincent et al. 2008). The implication is that individual-focused analyses can resolve details of regional specialization, particularly in association regions like the IFG, that may have densely packed functional zones that vary in location across individuals (Mueller et al. 2013).

The close juxtaposition of multiple functionally distinct regions near to language regions may also have complicated group-based functional connectivity estimates of network organization. Using datadriven algorithms that ‘parcellate’ the cortex into discrete networks, group-averaged analyses indicate the association cortices typically comprise around 5 major networks (Yeo et al. 2011; Power et al. 2011; Doucet et al. 2011), none of which is an unequivocal candidate for a language system (see Ji et al. 2019 for a discussion). Instead, other known distributed networks like the default (Buckner et al. 2008; see also Binder et al. 2009), frontoparietal control (Vincent et al. 2008) and salience networks (Seeley et al. 2007) have typically been identified within the vicinity of classical perisylvian language areas (e.g., see Fig. 11 in Yeo et al. 2011). The juxtaposition of language regions near to other major, dominant association networks may have obscured their identification in low-dimensional, low-resolution estimates of network organization. More recently, using network parcellation approaches, Gordon and colleagues (Fig. 3 in Gordon et al. 2017b; 2017a; Laumann et al. 2015) and Kong and colleagues (Fig. 2 in Kong et al. 2019) each delineated a network that matches the expected distribution of the language network. In these schemes, the network nearest to what might be a candidate language network was given the labels ‘ventral attention network’ and ‘temporal parietal’ network, respectively, highlighting the uncertainty over its function.

Analyses that assume a left-lateralized language network will be present yield clear positive evidence. For example, using a clustering approach that built in priors to nudge the algorithms to identify a distinct network anchored in the left superior temporal gyrus, Lee and colleagues identified a candidate language network (see LAN1 in Fig. 7 of Lee et al. 2012) that exhibited the hallmarks of the classic language system (i.e., containing regions in IFG, pSTC and TP). Interestingly, this network was found to also include smaller regions distributed along the midline in dorsal posteromedial cortex (dPMC), midcingulate cortex (MCC), ventromedial prefrontal cortex, and anterior superior frontal gyrus (aSFG). This ‘extended’ language network was recapitulated by Hacker and colleagues (2013), who used regions of activation from a metaanalysis of various cognitive domains to guide network definition (see Figs. 5 and 7 in Hacker et al. 2013; see also Hampson et al. 2002). Reinforcing the role of this extended network in language, Glasser et al. (2016; and see Ji et al. 2019) demonstrated that a similar network, defined through a multimodal approach, reveals task response during story listening. The identified regions fall at or near regions important to less domain-restricted aspects of cognitive control.

Thus, the complex literature on the network organization near to the frontal language regions almost certainly arises, in part, because there exist multiple distinct juxtaposed networks that are simply difficult to disambiguate in group-based studies. Fedorenko and colleagues’ (2010; 2012) findings within individuals of spatial separation of prefrontal language regions from adjacent domain-flexible processing regions provides a compelling demonstration that a more complete description of organization may be possible when fine anatomical details are preserved.

Motivated by the ambiguity of the prior literature and the opportunity to examine network organization fully within the individual, we sought to revisit and expand examination of the human language network. Specifically, we aimed to explore the detailed anatomy of the language network and contextualize it alongside other neighboring functional networks including the default, frontoparietal control and salience networks. What emerged is evidence that the language network is spatially distinct from but similarly organized to other differentially specialized association networks. Moreover, while sharing the same organizational motif as the other higher-order association networks, regions within the language network have particularly close spatial adjacencies to a network hierarchy involving motor and sensory regions important for speech and hearing.

## Methods

### Overview

The functional architecture of the language network of the cerebral cortex was explored using functional connectivity within individuals based on two approaches: manually selected seed-based connectivity and data-driven clustering. In all individuals tested, a clear candidate language network was observed that occupied regions juxtaposed but distinct from other distributed association networks including the default, frontoparietal control and salience networks. Next, data collected during a language localizer task were used to reveal language-responsive regions of the cortex. The candidate language network defined by functional connectivity overlapped in detail with regions activated by the task (Fedorenko et al. 2010). The analyses provided evidence that the extended language network, including smaller and previously underemphasized regions, responds to language task demands, supporting the idea that the distributed network is specialized for language. Data collected during a motor localizer task were used to define motor regions activating during tongue movements (n = 2 subjects) to explore the relationship between language network regions and sensorimotor cortices.

### Participants

Seven healthy adults (6 right-handed) were recruited from the greater Boston community and screened to exclude a history of neurological or psychiatric illness. Participants provided written informed consent using procedures approved by the Institutional Review Board of Harvard University. Data were collected as part of two separate studies. In Study 1, two subjects (2 females; ages 23 and 24) were each scanned across 24 separate MR sessions collected over approximately 16 weeks that included a language-localization task (resting-state data previously reported in Braga and Buckner 2017). Two additional potential subjects were excluded because of missing language task data and the absence of a fieldmap. In Study 2, five subjects (3 females; ages 20–25) were each scanned across 4 MR sessions collected over two weeks (portions of data previously reported in DiNicola et al. 2019). A sixth potential subject was excluded due to missed task trials during periods when the subject also had her eyes closed.

### MRI Data Acquisition

The detailed data acquisition protocol was previously reported in Braga and Buckner (2017) and DiNicola et al. (2019). Procedures are briefly summarized here. Data were acquired at the Harvard Center for Brain Science on a Siemens Prisma-fit 3T MRI scanner using the vendor’s 64-channel phased-array headneck coil (Siemens, Erlangen, Germany). Subjects provided behavioral responses during the language localizer task using a custom button box. Eyes were monitored and video-recorded using an Eyelink 1000 Core Plus with Long-Range Mount (SR Research, Ottawa, Ontario, Canada). A 4-point scale was used to record participant’s level of arousal during each run based on the frequency and duration of eye closures. The eye video was also visually checked to flag prolonged eye closures occurring during task trials.

Blood oxygenation level-dependent (BOLD) fMRI (Kwong et al. 1992; Ogawa et al. 1992) data were acquired using a multi-band gradient-echo echo-planar pulse sequence (Setsompop et al. 2012) implemented as part of the Human Connectome Project (HCP; Van Essen et al. 2013; Xu et al. 2012): TR 1000 ms, TE 32.6 ms, flip-angle 64°, 2.4 mm isotropic voxels, matrix 88 × 88 × 65, multi-slice 5× acceleration. Minimization of signal dropout was achieved by automatically (van der Kouwe et al. 2005) selecting a slice plane 25° from the anterior-posterior commissural plane towards the coronal plane (Weiskopf et al. 2006; Mennes et al. 2014). A rapid T1-weighted anatomical scan was acquired in each session using a multi-echo MPRAGE three-dimensional sequence (van der Kouwe et al. 2008): TR 2200 ms, TE 1.57, 3.39, 5.21, 7.03 ms, TI 1100ms, flip angle 7°, 1.2 mm isotropic voxels, matrix 192 × 192 × 144, in-plane GRAPPA acceleration 4. A dual-gradient-echo B0 fieldmap was acquired to correct for spatial distortions: TE 4.45, 6.91 ms with matched slice prescription/spatial resolution to the BOLD sequence.

Functional runs were flagged for exclusion if 1) maximum absolute motion exceeded 2mm, 2) slice-based temporal signal-to-noise ratio was less than or equal to 135, or 3) the value for maximum absolute motion or signal-to-noise ratio represented an outlier when values from all runs were plotted together. The raw data from flagged runs were visually checked for motion artifacts and excluded if these were deemed to be severe. All exclusions were determined prior to analysis of the task data. Following this procedure, one language-localizer run was excluded for S6 due to high motion and low SNR. Furthermore, 4 out of 24 fixation runs and 1 out of 8 language localizer runs were excluded for S2 after detection of signal instability (higher mean signal compared to other runs) that was later determined to arise from the gradient coil.

### In-Scanner Tasks

All 7 participants provided data collected during multiple runs of a passive visual fixation task and a language localizer task (Fedorenko et al. 2010). Study 1 also included multiple runs of a motor localizer task. Table 1 outlines the number of BOLD runs collected and included from each participant for each task. For all tasks, stimuli were projected onto a screen located behind the participants’ head and viewed through a mirror. Participants were instructed to remain still, stay awake, and stay engaged for the duration of each run. Both studies included additional tasks that were not analyzed here.

**Table 1.** Number of runs included/collected from each subject.

| Subject | Study | Fixation | Language Localizer | Motor Localizer |
| --- | --- | --- | --- | --- |
| S1 | 1 | 24/24 | 8/8 | 8/8 |
| S2 | 1 | 20/24 | 7/8 | 8/8 |
| S3 | 2 | 8/8 | 8/8 | - |
| S4 | 2 | 8/8 | 8/8 | - |
| S5 | 2 | 8/8 | 8/8 | - |
| S6 | 2 | 8/8 | 7/8 | - |
| S7 | 2 | 8/8 | 8/8 | - |

Notes: Study 1 included a fixation, a language localizer and a motor localizer task. The numbers below each task label indicate the number of runs included / collected. Study 2 included a fixation and a language localizer task only. Individual runs were excluded based on criteria that included head motion and signal-to-noise ratio thresholds, as well as sleepiness in the scanner.

#### Fixation Task

The fixation data were used for functional connectivity definition of networks. Participants fixated a black ‘+’ symbol presented at the center of a light gray screen. Each run lasted 7m 2s. In Study 1, data were collected over 24 MR sessions each of which included one fixation run (total 168m 48s of fixation data per individual). In Study 2, data were collected over 4 MR sessions, each of which included two fixation runs (total 56m 16s of fixation data per individual). Fixation data from Study 1 participants (S1 and S2) were previously reported in Braga and Buckner (2017; referred to as ‘S4’ and ‘S3’, respectively, in that study) but were preprocessed differently here using updated strategies to minimize spatial distortion and blurring (employed in Braga et al. 2019)

#### Language Task Contrast

Participants performed 8 runs of the language localizer task developed by Fedorenko et al. (2010). This task contrasted reading meaningful sentences versus lists of pronounceable non-words. Basic task requirements were matched between conditions (e.g., engaging with visual stimuli with the same visual features, performing button presses, and phonological processing), while preserving lexico-semantic and syntactic processing in the sentence condition (Fedorenko et al. 2010). The language localizer task contrast reveals activation in classical language regions of the IFG, pSTC and TP, as well as the cerebellum. Furthermore, the regions show spatial specificity in relation to juxtaposed regions as well as functional specificity in relation to non-linguistic task demands (see Fig. 2 in Fedorenko et al. 2011; Fedorenko et al. 2010; 2012). Both visual and auditory versions of the task identify a similar set of regions (Braze et al. 2011; Scott et al. 2016), supporting that the regions are responsive to language independent of input modality.

Participants fixated a black ‘+’ symbol on a light gray background and read sentences (S; e.g., ‘TOM GOT MARRIED TO A LAWYER LAST YEAR AND SEEMED VERY HAPPY’) and lists of pronounceable non-words (N; e.g., ‘CRE ENFENTLY SILE U ALGOW OLP LENSIS ZOLLER NALD LIRM U LAS’). Sentences were presented centrally on the screen one word at a time. Each sentence was composed of 12 words, with each word presented for approximately 0.45 s. Each sentence was followed by a cue (an image of a finger pressing a button) lasting 0.50 s that instructed participants to press a button with their right index finger. The button response was included to keep participants engaged. Each sentence lasted ~6.2 s.

The task began with an 18 s fixation period (+). A blocked design was used, where each condition (S or N) was presented as a block containing three consecutive trials. Four alternating blocks were presented sequentially, followed by a fixation period lasting ~15.6 s. Twelve blocks were presented in each run. The condition order was counterbalanced across two different designs that were each performed 4 times by each participant, leading to 8 runs collected from each participant. The designs were: 1; +, S, N, S, N, +, N, S, N, S, +, S, N, S, N, +; and 2; +, N, S, N, S, +, S, N, S, N, +, N, S, N, S, +. For the targeted contrast, the sentence conditions were contrasted with the non-words condition (S > N; see *Task Activation Analyses*). The total task duration was 300 s for each run (total: 40m of language localizer data per individual). Every sentence across all sessions within an individual was unique.

#### Motor Task Contrasts

To estimate regions active during tongue movements, participants performed a series of subtle controlled movements in the scanner (adapted from Buckner et al. 2011). Participants were trained prior to scanning on three types of movements: finger taps (sequentially touching the index and middle fingers to the thumb), foot taps (subtle dorsiflexion and plantar flexion), and tongue movements (touching the canines on the left and right side with the tip of the tongue with lips closed). Movements were made in a way that minimized muscle tension and movement of other body parts. A blocked design was employed with 5 active task conditions: left hand (LH), right hand (RH), left foot (LF), right foot (RF), and tongue (T) movements. Passive fixation (+) occurred between active conditions and also began and ended the run.

Each condition lasted 18s, during which a cue stimulus (an illustration of the relevant body part) and words describing the condition (e.g., ‘LEFT HAND’) were shown. The cue flickered on and off on a 1-Hz cycle, and participants were cued to make the movements to the timing of the flicker. An index and a pointer finger movement were performed per cycle in the hand condition, one foot tap (dorsiflexion and plantar flexion) was performed per cycle in the foot condition, and a left and right movement were performed per cycle in the tongue condition. Participants were visually monitored while they performed these movements by the scanner operators to ensure compliance. The condition order was counterbalanced across two different designs that were each performed 4 times, leading to 8 runs per participant. The two designs were: 1; +, LH, RH, T, LF, RF, +, RF, LF, T, RH, LH, +; and 2; +, LF, RF, T, LH, RH, +, RH, LH, T, RF, LF, +. The targeted contrast was intended to isolate the orofacial motor region, hence the tongue movement condition was contrasted with all other conditions (i.e., T > LH + RH + LF + RF; see *Task Activation Analyses*). The total task duration was 234 s for each run (total: 31m 12s motor localizer data per subject).

### MRI Data Processing

#### Within-Subject Data Alignment

Data processing procedures were previously described in detail in Braga et al. (2019) and are summarized here. An inhouse pipeline (‘iProc’) optimized alignment of within-subject data collected across different scanning sessions, preserving anatomical detail as much as possible by minimizing spatial blurring and multiple interpolations (expanding on Braga and Buckner 2017; Yeo et al. 2011; Poldrack et al. 2015). Each subject’s data were processed separately. To optimize alignment, two subject-specific registration templates were created: a mean BOLD template and a T1 native-space template. BOLD data from every run contributed to the subject’s mean BOLD template, minimizing bias towards any run or session. The T1 native-space template was created by selecting a T1-weighted structural image (upsampled to 1mm isotropic space) that was visually deemed to have good pial and white matter boundary surface estimates (see *Projection to Cortical Surface*).

For each BOLD volume, three transforms were calculated to 1) correct for head motion, 2) correct for geometric distortions caused by susceptibility differences using a B0 fieldmap, and 3) register the BOLD volume to the within-subject mean BOLD template. A further transform was calculated once for each subject and applied to all registered volumes which projected data from the mean BOLD template to the T1 native-space template. The four transformation matrices were composed into a single matrix that was applied to each original BOLD volume to project all data to the T1 native space-template in a single interpolation. The iProc pipeline yielded data aligned to a subject-specific template at 1-mm isotropic resolution, with minimal interpolation and signal loss.

#### Additional Processing for Functional Connectivity

For functional connectivity analyses, additional processing steps included regression of nuisance variables and bandpass filtering. Nuisance variables included 6 motion parameters plus whole-brain, ventricular and deep white matter signal, and their temporal derivatives. These signals were regressed out of native-space-projected BOLD data (using 3dTproject; AFNI v2016.09.04.1341; Cox 1996; 2012). This was followed by bandpass filtering at 0.01–0.1 Hz (using 3dBandpass; AFNI v2016.09.04.1341; Cox 1996; 2012).

#### Projection to Cortical Surface

Pial and white matter boundaries were calculated automatically using FreeSurfer’s recon-all (Fischl et al. 1999). Data were resampled from the native space to the fsaverage6 standardized cortical surface mesh (containing 40,962 vertices per hemisphere; using mri_vol2surf; Fischl et al. 1999) and then surface-smoothed using a 2mm FWHM kernel. Data were sampled from the gray matter at a position halfway between the white and pial surfaces using trilinear interpolation. For task-based analyses, BOLD data prior to nuisance regression and bandpass filtering were projected.

### Functional Connectivity Analyses

Functional connectivity analyses were performed on the surface using both seed-based and unbiased data-driven parcellation techniques. For the seedbased approach, pair-wise Pearson’s product-moment correlations between the fMRI timeseries at each vertex were computed, yielding an 81,924 × 81,924 correlation matrix (40,962 vertices per hemisphere) for each run of BOLD data. These matrices were Fisher-transformed and averaged together yielding a within-subject across-run mean correlation matrix with high stability for each subject. This average matrix was then inverse-Fisher-transformed back to correlation values and assigned to the vertices of a cortical template created in-house (as described in Braga and Buckner 2017). This template allowed individual vertices to be selected for real-time visualization of the resulting correlation maps using the Connectome Workbench’s wb_view software (Marcus et al. 2011). For final visualization of seed-based connectivity maps, correlation values were converted back to z(r) using the Fisher-transform.

#### Initial Observation and Hypothesis

The observation that a candidate language network may be detectable within individuals was made during a previous exploration of the functional anatomy of the default network in S1 (referred to as ‘S4’ in Braga and Buckner 2017). While manually selecting seeds, a distinct network was observed that followed the distributed motif of other association networks, but occupied separate regions of the cortex. Notably the network contained large regions in the left lateral temporal and left inferior frontal cortices, near classical language areas. The hypotheses were formed that i) the anatomical details of this candidate language network could be reproducibly defined in additional subjects, and ii) the network would show increased activity during a task targeting linguistic processes. We targeted these hypotheses using network mapping techniques and by comparing the network maps with regions activated during a language localizer task (Fedorenko et al. 2010). Critically, while the initial discovery was made using manual procedures, the observations were converged upon by automated classification.

#### Manual Targeting of Candidate Language Network

To identify the candidate language network in additional subjects, seed regions were manually selected from the left prefrontal cortex, at or near to where the precentral sulcus meets the posterior middle frontal gyrus (i.e., pMFG; Fig. 1). This region was targeted because it contained a prominent region of the candidate language network in the initial exploration of S1 (see also Glasser et al. 2016; Fedorenko et al. 2010; Lee et al. 2012). An iterative process was used for seed selection, similar to that described in Braga and Buckner (2017) and Braga et al. (2019). A seed vertex was identified in each individual that revealed a robust network with a spatial distribution that resembled the candidate language network as initially observed in S1. Correlation maps were thresolded at z(r) > 0.2 for visualization and displayed with the Jet look-up-table (colorbar) set to a range between 0.2–0.6. A network was deemed robust if it generally revealed high correlation values (z(r) ≈ 0.6), but also if the network regions displayed sharp boundaries (surrounded by areas of low correlation). Specifically, to assure that the candidate language network was being detected selectively, the observer’s knowledge of spatial features from other networks was also used in seed selection. For example, candidate seed vertices were not selected if they revealed prominent connectivity to the posterior midline at or near the cingulate and retrosplenial cortices, which are hallmark features of the default network. Similarly, candidate seed vertices revealing patterns resembling the frontoparietal control network were not selected. In other words, the seed-selection process targeted specific features of the initially observed candidate language network and excluded features of other known networks.

**Figure 1:**
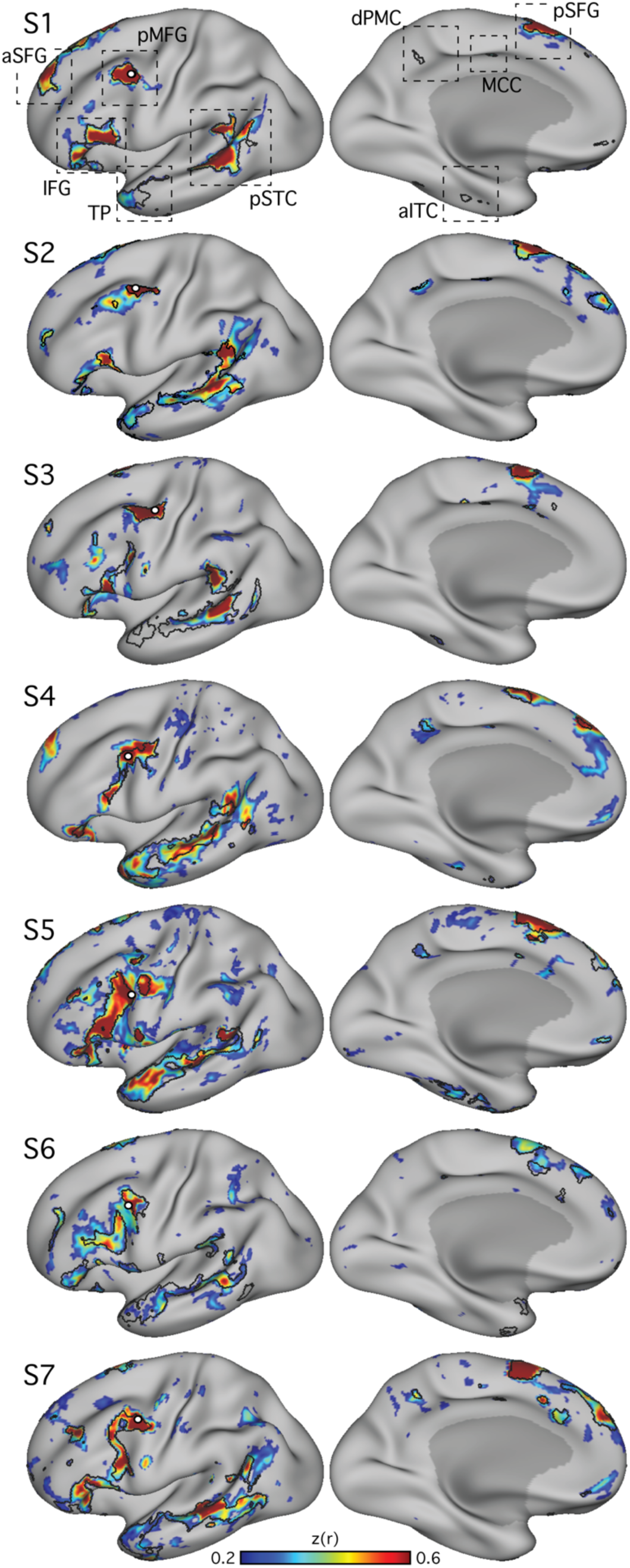
Within-individual intrinsic functional connectivity identifies a candidate distributed language network. Seven subjects (S1–S7) each reveal a candidate language network. Seed regions (small white circles) are displayed at or near the posterior middle frontal gyrus (pMFG). Correlation patterns are shown on an inflated cortical surface representation of the left hemisphere. In each subject, the correlation patterns (colorbar) show a network that included regions located near to classical language regions of the inferior frontal gyrus (IFG; Broca’s area) and posterior superior temporal cortex (pSTC; Wernicke’s area). The network also revealed regions distributed across multiple cortical zones (see dashed boxes in top panel) including the posterior superior frontal gyrus (pSFG), the anterior superior frontal gyrus (aSFG; appearing in medial and/or lateral portions in different subjects), and the temporal pole (TP). Smaller regions observed consistently in 5 or more subjects included the dorsal posterior medial cortex (dPMC), the middle cingulate cortex (MCC), and the anterior inferior temporal cortex (aITC). Lateral (left column) and medial (right column) views are shown. z(r), Fisher’s r-to-z transformed Pearson’s product-moment correlations.

#### Confirmation of the Network from Distributed Cortical Zones

To determine if the network was spatially selective and similar if defined outside of prefrontal cortex, additional seed regions were examined in two subjects (S1 and S2). The approximate locations of regions revealed by the original pMFG seeds were targeted in the anterior and posterior IFG, the pSTC, and the posterior superior frontal gyrus (pSFG; Fig. 2). In each zone, for each subject, the iterative seed selection process was again followed, resulting in a single seed that targeted the candidate language network in each cortical zone and subject. A similar network was detectable from seeds in all zones.

**Figure 2:**
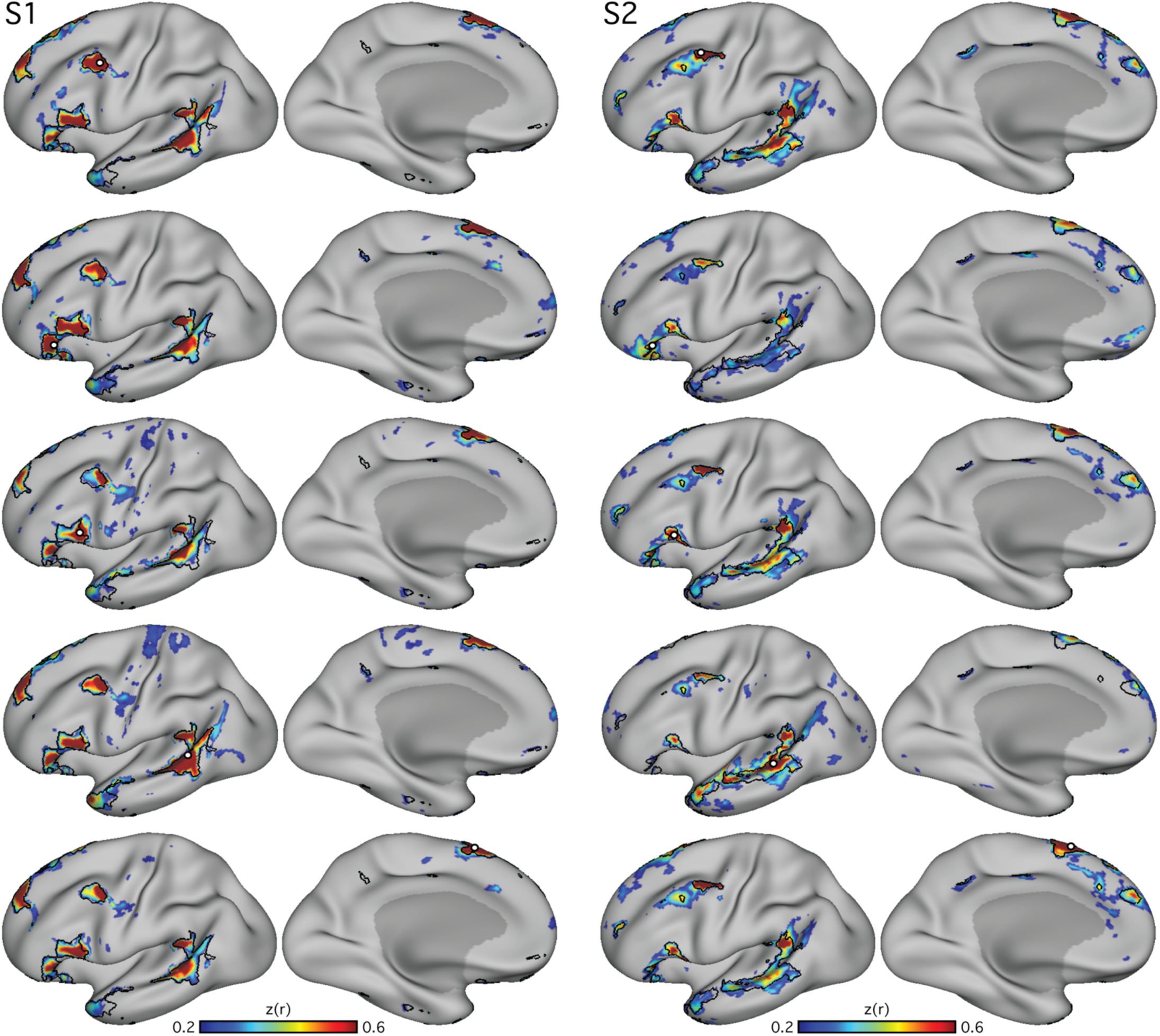
Distributed organization of the candidate language network is confirmed using seed regions in multiple cortical locations. In two subjects (S1 and S2), seed regions (small white circles) were selected from different portions of the network identified in Fig. 1. In each panel, the candidate language network defined by data-driven parcellation (see Fig. 4) is shown in black outline, to provide landmarks for comparing across panels. In each subject, seed regions were placed in the inferior frontal gyrus at an anterior (second row from top) and posterior site (third row), as well as in the posterior superior temporal sulcus (fourth row) and posterior superior frontal gyrus (last row). Although the maps differ in their details, the large-scale distribution and location of the network regions are appreciably similar across seed regions, with regions of high correlation falling generally within the parcellation-defined boundaries. z(r), Fisher’s r-toz transformed Pearson’s product-moment correlations.

#### Generalization of the Candidate Language Network Across Acquisition Task States

To explore whether the detection of the candidate language network was dependent on the behavioral state of participants during data acquisition, functional connectivity was performed using data acquired during three different tasks. Data from the fixation, language localizer and motor localizer tasks were analyzed separately for two subjects (S1 and S2). For each task, in each subject, initially the same seed vertex as previously selected from the fixation data (see *Manual Targeting of Candidate Language Network*) was used. If this seed failed to produce a robust map in the other two task datasets, another seed was selected at or near the pMFG following the iterative process described above. A similar network was detectable using data from all three tasks (Fig. 3).

**Figure 3:**
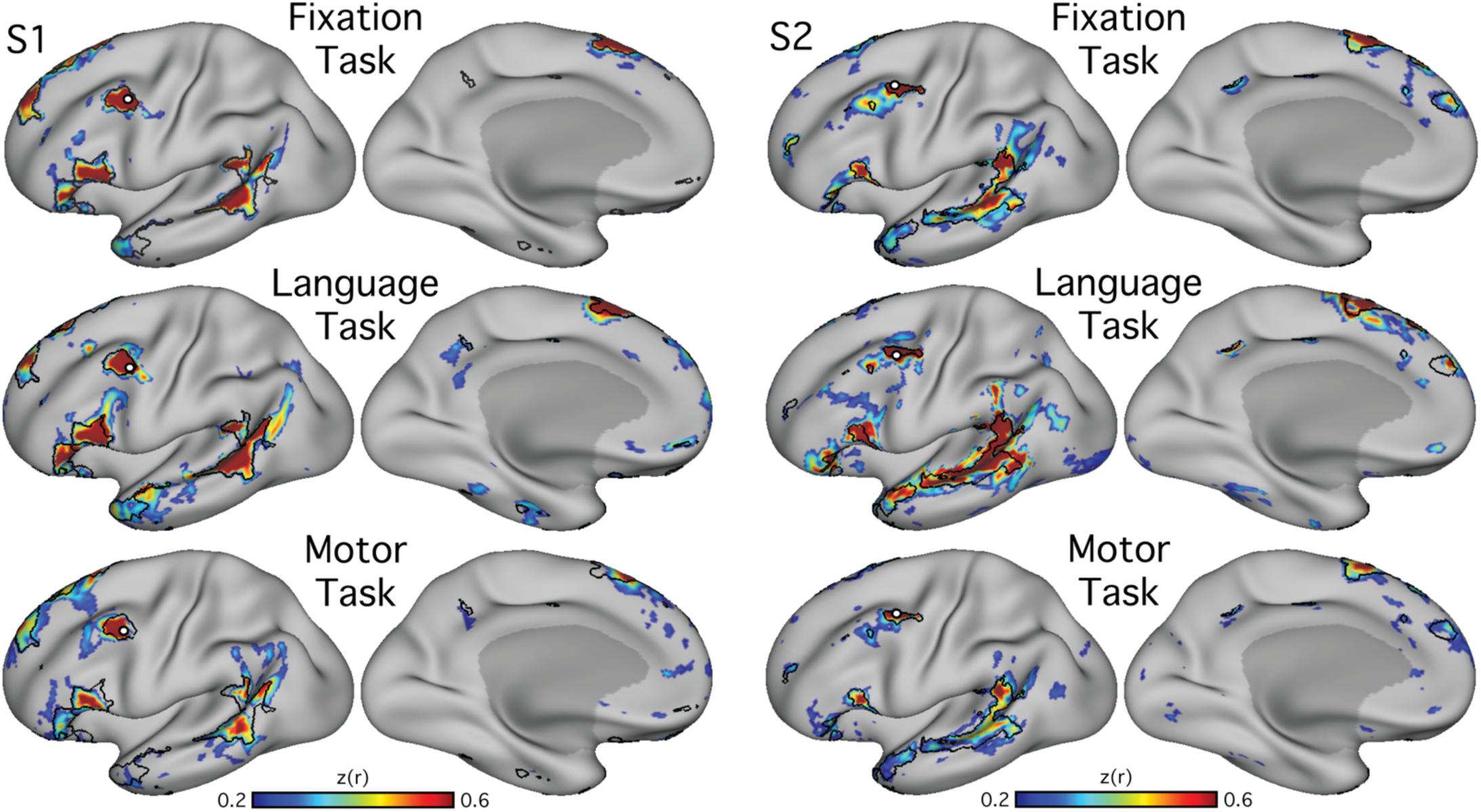
The connectivity-defined candidate language network generalizes across data acquired in different task states. Functional connectivity reliably defined the candidate language network across three distinct tasks, showing that the presence of the network is not dependent on a specific cognitive context (see text for task descriptions). Note that the location of the seed region (small white circles) was optimized for each data set to show that the topography of the network is stable despite minor differences in functional shifts that might occur due to task context. Note that the optimal seed location also varies across data sets even when collected during the same task context (see Supplementary Figure S3 in Braga and Buckner 2017, and Figure 3 in Braga et al. 2019). z(r), Fisher’s r-to-z transformed Pearson’s product-moment correlations.

#### Data-driven Parcellation

Although seed-based correlation is able to reveal the topography of the intrinsic networks, it relies heavily on observer input, which could result in bias. To confirm that the definition of the candidate language network was not a consequence of observer bias, a data-driven parcellation analysis was performed for each subject using *k*-means clustering. Preprocessed BOLD data from the fixation task were concatenated in time and MATLAB’s *kmeans* function (v2015b; MathWorks, Natick, MA) was used to cluster the timeseries into networks. Default settings were used (1 random initialization, 100 iterations, squared Euclidean distance metric). As the results will reveal (Fig. 4), similar network estimates were found for both the seed-based and parcellation approaches suggesting that the identification of the candidate language network is robust to different network discovery methods. *K*-means clustering was performed in each individual at *k* = 17 (as in Yeo et al. 2011). It is important to note that the clustering and seed-based approaches yield similar but not identical network estimates, and that the network topography is influenced by the number of clusters defined. In the present analyses, *k* = 17 was used because it produced a network that was found to correspond to the candidate language network as defined by seed-based connectivity, and it recapitulated other previously observed distinctions between distributed networks (Braga and Buckner 2017), in all individuals.

**Figure 4:**
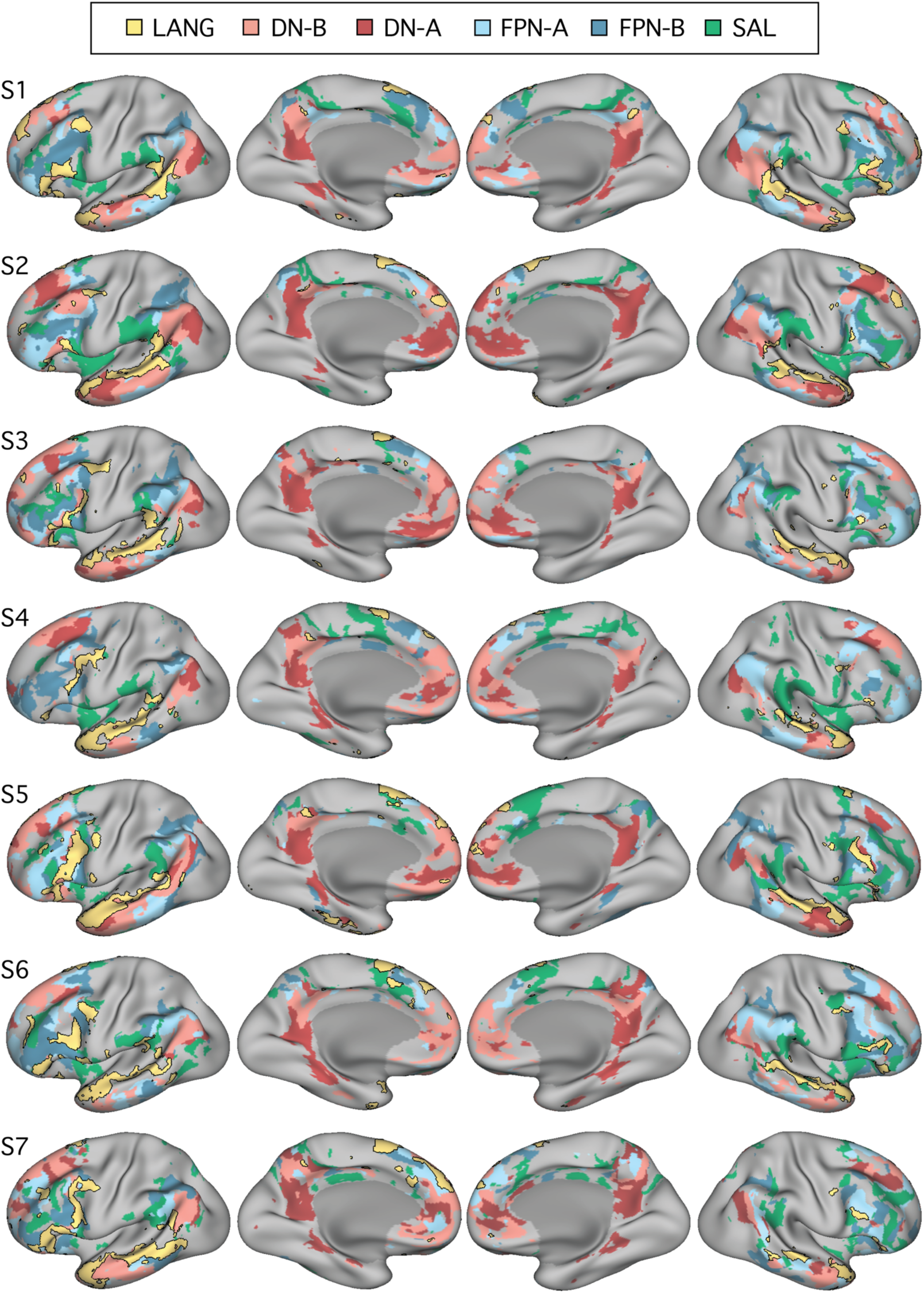
Close juxtaposition of the candidate language network with neighboring distributed networks revealed by data-driven parcellation. K-means clustering was used to parcellate the cortex into 17 discrete networks. The candidate language network (LANG; yellow and black outline) was observed in all participants (S1–S7). Network regions were recapitulated in all of the nine zones highlighted in Fig. 1, including a region in the temporal pole that extended rostrally. Further regions can also be observed in the right hemisphere. From the parcellation solutions, five additional networks were selected for further analysis due to their spatial proximity to the language network and their identification within classic language regions in prior data-driven network analyses (e.g., Yeo et al. 2011). These networks were the salience network (SAL; green), frontoparietal control network-A and -B (FPN-A and FPN-B; blues), and default network-A and -B (DNA and DN-B; reds). The LANG network had a complex spatial relationship with these neighboring networks, showing regions closely packed with default, frontoparietal control and salience network regions in the temporal cortex, and inferior and dorsal frontal cortices. The left two columns show lateral and medial views of the inflated left hemisphere, while the right two columns show the right hemisphere.

#### A Priori Selection of Networks

In order to explore language-driven task responses in relation to the spatial distributions of multiple closely juxtaposed association networks, 5 networks, in addition to the candidate language (LANG) network, were selected for further analysis from the 17-network parcellation. The selected networks included: the two networks previously identified within the canonical default network (DN-A and DN-B), two networks that are positioned near to the canonical frontoparietal control network (FPN-A and FPN-B; see Braga and Buckner 2017), and the salience network (SAL; Seeley et al. 2007; Dosenbach et al. 2007). The networks were identified and labeled according to previously described anatomical features (Braga and Buckner 2017; Dosenbach et al. 2007). Anatomical details of FPN-A and FPN-B were previously reported for two subjects (including subject S1, labeled ‘S4’ in Braga and Buckner 2017).

As can be seen in Fig. 4, the networks differed in their detailed anatomy across subjects. Specific spatial relationships served as useful anchoring points but, given the complex relationships of the networks, any assignment must be considered a hypothesis awaiting independent functional confirmation to build confidence (such as provided for DN-A and DNB in DiNicola et al. 2019 and sought here for the LANG network). That said, certain features and patterns are largely consistent across subjects. FPN-A and FPN-B both occupy regions of the lateral inferior frontal cortex and parietal regions at or near the intraparietal sulcus. Within the inferior parietal lobule, FPN-A typically occupies a region more ventral to FPN-B, and more anterior to DN-B. Even so, these regions are heterogeneous and difficult to match across subjects. Perhaps the most reliable identifying feature is that the DN-A, DN-B, FPN-A and FPN-B networks follow a stereotyped anterior-posterior sequence along the inferior lateral temporal cortex. The relative position in this portion of the brain of FPN-A (anterior) and FPNB (posterior) served as a useful guide for labeling those networks. In all subjects, one of the 17 networks defined by clustering was deemed to correspond with each of the DN-A, DN-B, FPN-A and FPN-B networks based on these previously reported anatomical features.

The SAL network was identified by the presence of regions in the anterior inferior parietal lobule or supramarginal gyrus, in the inferior frontal cortex and insula, and a region or set of regions along the dorsal midline, sometimes circling the medial somatomotor cortex in a ‘U’ shape (Fig. 4). The similar large-scale distribution of the SAL network regions across subjects offers some confidence that the same broad network was being targeted. For example, note that the parietal region of the SAL network was located in the supramarginal gyrus, anterior to FPN-A, in all subjects. However, the correspondence was not perfect, with gaps evident between the SAL network parietal region and the other network regions in some subjects. In each subject, the network that most closely followed the anatomy of the canonical SAL network was chosen.

In this way, 5 additional distributed association networks were identified *a priori* that were all near to the LANG network regions. These networks were each tested for task-driven response during the language localizer task contrasts.

### Task Activation Analyses

Data were analyzed for task-driven response using the general linear model as implemented by FSL’s FEAT (Woolrich et al. 2001). Preprocessed and smoothed data from each BOLD run were entered into a first-level analysis. Surface-projected data from the left and right hemispheres were analyzed separately, and the results were combined after for visualization and *a priori*-defined network activation analysis. The data and model were highpass filtered using a cut-off of 100s to reduce the influence of low frequency noise. A linear term was included in the model to account for linear drifts in the data. Each task condition was modelled as a separate explanatory variable using a block design (see *In-Scanner Tasks*). The explanatory variables were convolved with a double-gamma haemodynamic response function. Temporal derivatives were included in the model to account for variations in the haemodynamic response. In the language localizer task, for the targeted contrast of sentences > nonwords conditions, at each vertex the beta value for the non-word condition was subtracted from the beta value for the sentences condition. In the motor localizer task, for the contrast of tongue movements > other movements, at each vertex the beta value for the tongue condition was multiplied by 4, and the sum of the beta values for the right hand, left hand, right foot and left foot conditions were subtracted. For both contrasts, the resulting values at each vertex were converted to *t* statistics by dividing by their standard error, and then converted to a *z*-statistic. Within each subject and task, the *z-*statistic maps from all runs were averaged together using *fslmaths* (Smith et al. 2004).

For visualization, *z* thresholds were selected to best demonstrate the task activation patterns for each subject (thresholds: S1, 3.0–8.0; S2, 3.5–8.0; S3, 3.5– 8.0; S4, 5.0–14.0; S5, 3.5–10.0; S6, 3.0–7.0; S7, 2.0–6.0; Fig. 5). A lower threshold was picked just above that needed to remove low-confidence activations (i.e., small, randomly dispersed spots or speckles showing low *z* values), and an upper threshold was picked that allowed vertices of low and high correlation within the contiguous regions to be discerned.

**Figure 5:**
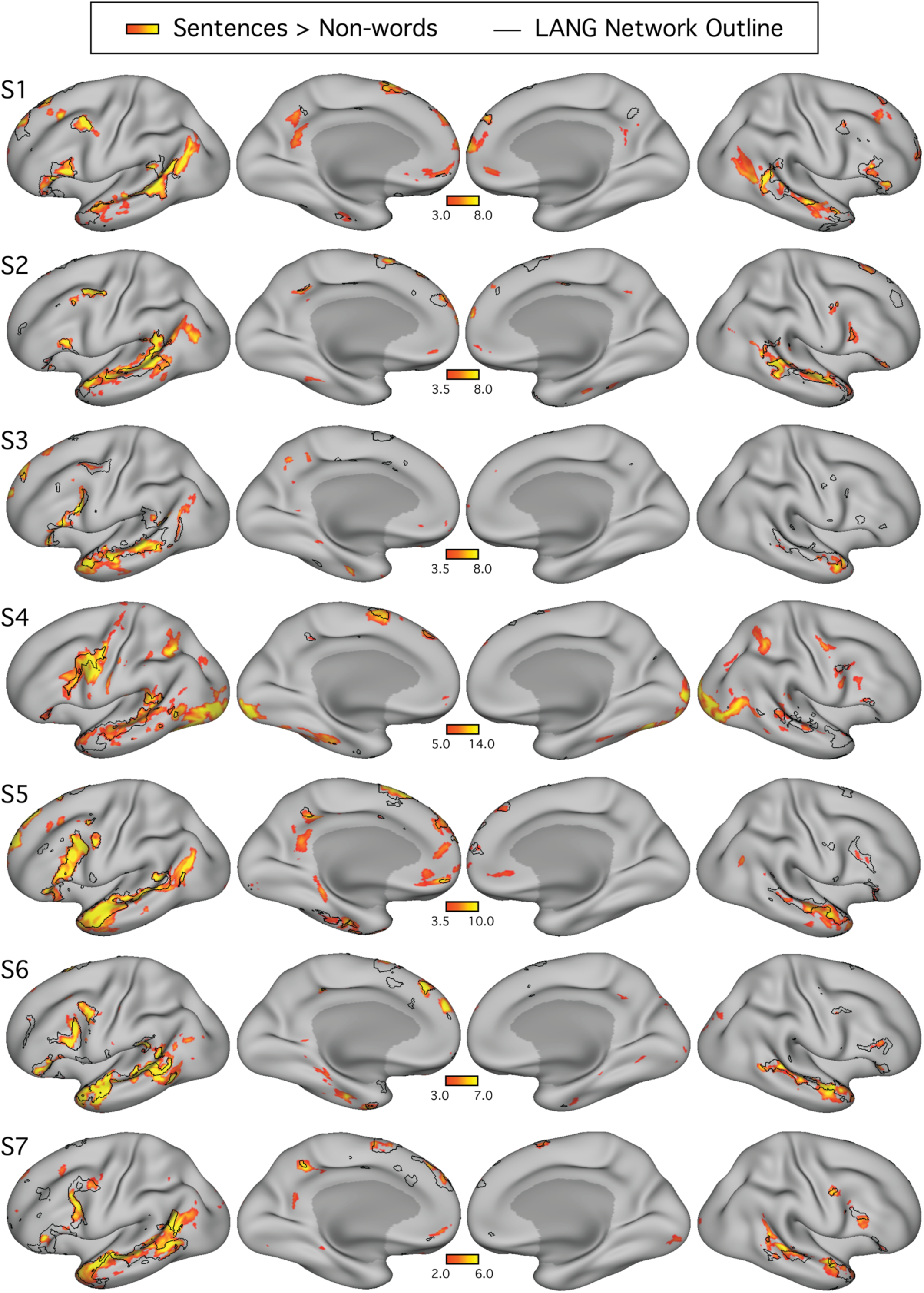
The candidate language network shows close spatial correspondence with regions activated during a language task contrast. The language network (LANG) is shown in black outline and was defined using k-means clustering. Independently acquired data collected during a language localizer task contrast (Fedorenko et al. 2010) reveals cortical response to linguistic demands. Red-yellow colorbars show within-individual z-normalized beta values (i.e., ‘increased activation’) for the contrast of reading sentences versus reading lists of non-words. In all subjects (S1–S7), the language task activations fell largely within the boundaries of the intrinsically defined candidate language network. The overlap was not perfect, and in some cases hints of other networks can be seen (e.g., see S4 and S5), though these exceptions were not consistent across subjects. The upper and lower thresholds were selected by eye for each subject, to show the distribution of language-responsive regions while removing regions showing low responses. The detailed anatomy of the distributed intrinsic network corresponds closely with regions showing task-driven activation, including in smaller areas extending beyond the classical language zones (e.g., see S2 and S6), suggesting that the intrinsically organized network is functionally specialized.

A key question was whether the topography of the task contrast map for the language localizer task corresponded to the topography of the intrinsic connectivity LANG network. To address this question, two approaches were used. First, the maps were visually compared: the spatial map from the parcellation analysis was overlaid onto the cross-run average task activation map (Fig. 5). Second, a network-of interest approach was used using the 6 *a priori* selected networks defined in each subject (see *A Priori Selection of Networks*). The average beta value for the contrast of sentences > non-words was calculated for all vertices falling within each network. Values from both the left and right hemispheres were included. Average beta values were calculated for each run of the language localizer task, leading to 8 estimates of the network’s recruitment during the task for each network and subject (except for S2 and S6 who each provided 7 runs; Table 1). The cross-run average beta value for each network was then plotted in a bar graph, along with the standard error of the mean (Fig. 6). This latter analysis has the benefit that there are no thresholds or subjective steps – the magnitude and variance of the response in each data-driven *a priori*-defined network is obtained and quantified in each individual.

**Figure 6:**
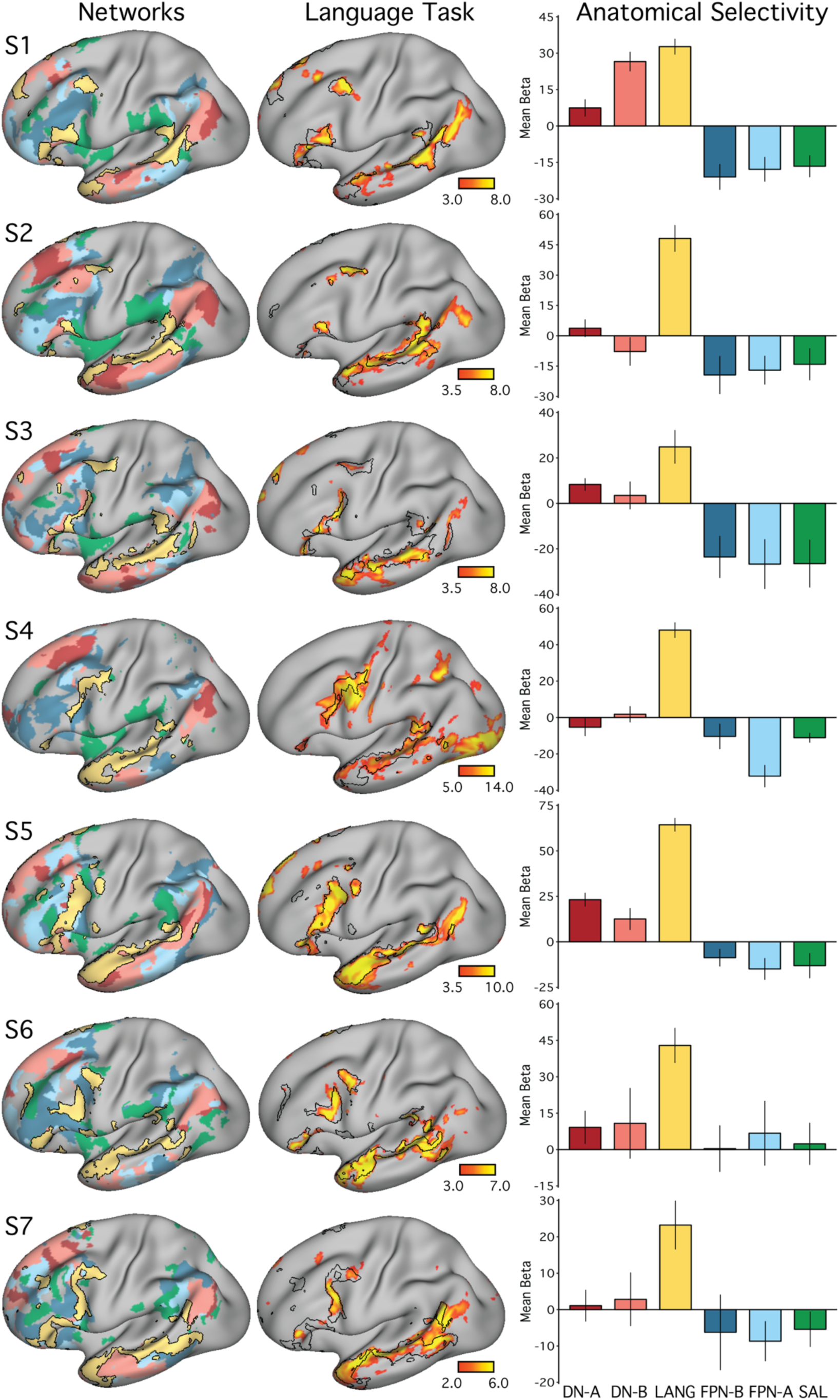
The candidate language network is selectively activated during a language task contrast. (Left column) The networks defined by intrinsic functional connectivity from Fig. 4 are replotted. The candidate language network (LANG) is shown in yellow, with the salience network (SAL) in green, the frontoparietal control networks (FPN-A and FPN-B) in blues, and the default networks (DN-A and DN-B) in reds. (Middle column) Task activation for the contrast of reading sentences versus reading lists of non-words (sentences > nonwords) is shown, with the intrinsic LANG network outline in black (see Fig. 5 for other views). (Right column) Bar graphs show the mean beta values for the sentences > nonwords contrast, averaged within each within-individual a priori-defined network, along with the standard error of the mean. Despite differences across individuals, LANG was the only network showing consistently higher activation for sentences > non-words, showed the highest activation of all networks in all participants, and in some cases (S2, S4 and S7) was the only network that showed clear increased activity for language.

### Experimental Design and Statistical Analysis

This study includes n = 7 participants, two of which were scanned over 24 brief MRI sessions and five of which were scanned across 4 extended sessions. All analyses focused on within-individual quantities. In all analyses, data were averaged over all usable runs that were collected from each individual (see Table 1). Functional connectivity between brain regions was calculated in MATLAB (version 2015b; http://www.mathworks.com; MathWorks, Natick, MA) using Pearson’s product moment correlations and Fisher’s r-to-z transformation prior to averaging across runs. Network parcellation was performed using MATLAB’s *kmeans* function (version R2015b). Task data were analyzed using the general linear model as implemented using FSL’s first-level FEAT (Woolrich et al. 2001). The cross-run average task activation map was created by taking beta maps from each run, *z*-normalizing and then averaging together using *fslmaths* (Smith et al. 2004).

## Results

### A Candidate Language Network is Identified by Functional Connectivity Within the Individual

The language network (LANG) was defined in all 7 individuals tested (Fig. 1) using seeds manually placed in the pMFG. In all cases a distributed network was observed that contained regions within the IFG, the pSTC, the TP, and the pSFG. The pSTC region sometimes extended into the inferior parietal lobule near to the supramarginal gyrus, but a clear and robust region in angular gyrus was not observed (see Figs. 1, 2 and 4). The LANG network contained further regions, extending to upwards of 9 cortical zones in the left hemisphere (highlighted in Fig. 1) replicating the extended language network defined by Lee et al. (2012) and Hacker et al. (2013; see also Glasser et al. 2016). A distinct region in the left anterior superior frontal gyrus (aSFG; appearing in medial and/or lateral portions in different subjects) was observed in all subjects. Regions in the dorsal posteromedial cortex (dPMC; at or near the posterior cingulate and precuneus), the middle cingulate cortex (MCC), and the anterior inferior temporal cortex (aITC) were observed in 5 subjects. In 4 subjects (S1, S4, S5 and S7), suggestion of a further region was observed at or near the ventromedial prefrontal cortex, despite this region suffering from signal dropout. The presence of a network region in each of the 9 highlighted zones in Fig. 1, replicated across a majority of individuals, suggests that the candidate language network is widely distributed and extends beyond regions that define the classical language system.

### The Candidate Language Network Generalizes Across Datasets and Analysis Methods

To support that the identified regions formed a distributed interconnected network, seeds were placed in 4 of the other large regions of the LANG network. In each case, the seeds produced correlation maps that were similar to that defined by the original pMFG seed (Fig. 2), suggesting definition of the LANG network was not dependent on a single seed location or vertex.

A further analysis tested whether the definition of the LANG network was dependent on the specific task that was performed during data acquisition. To address this question, data were analyzed from the same individuals during the performance of two additional tasks: the language and motor localizer tasks. In both cases, intrinsic connectivity from a seed in the pMFG revealed a similar distribution of regions as that identified using the visual fixation task data (Fig. 3). Subtle differences were observed. For instance, the correlations were generally higher, and the defined regions slightly larger, during the language task in S2. Similarly, in S1, the LANG region in the TP was emphasized in the language task data compared to the other tasks, and the pSTC region extended further into the angular gyrus. These differences could be a consequence of larger signal fluctuations being driven by the language task. Despite these differences, the same general distribution of regions was revealed across the three task contexts, including the active motor tasks.

The final analysis ensured that the definition of the LANG network was not a result of observer bias in the selection of seed regions. A data-driven parcellation approach to defining the networks (*k*means clustering) was performed. In all participants, parcellation revealed a candidate language network (Fig. 4) with near complete overlap with the network as defined by seed-based connectivity (see black outlines in Fig. 1) including smaller distributed regions (Figs. 1–3, see especially S1, S3 and S7 in Fig. 1).

An interesting difference was that in the temporal pole the clustering approach revealed a large region that was diminished or absent in the thresholded seed-based maps. The temporal pole suffers from signal dropout in MRI due to magnetic susceptibility differences with the nearby sinuses. It is possible that the parcellation approach is able to detect networks in regions of low signal because it clusters all vertices based on their relative pattern of correlations, rather than using an absolute correlation threshold.

### The Candidate Language Network is Bilateral but Left-Lateralized

In addition to the left-hemisphere regions detailed above, the LANG network also displayed multiple distinct regions in the right hemisphere (Fig. 4). The locations of these regions were roughly homologous to the zones observed in the left hemisphere, with a similar distributed organization including the right pMFG, IFG, pSTC, pSFG and TP in all subjects. Both hemispheres contained a large region spanning almost the length of the superior temporal sulcus. However, for other regions the right hemisphere homologs were visibly smaller in surface area (Fig. 4). In zones where evidence was found for small regions in the left hemisphere (pPMC, MCC, aITC), the homologous right-hemisphere regions were sometimes not observed.

It is important to note that the parcellation approach simultaneously clusters all surface vertices across both hemispheres. Hence the apparent leftright asymmetry in size observed in the clustering solution likely reflects actual differences in the network topology, as opposed to a spatial bias that can occur in seed-based approaches by selecting seeds from the left hemisphere. As a confirmation that the observed asymmetry was not a result of such bias, when seed regions were placed in the right pSTC region (biasing the correlations towards the right hemisphere) in some subjects, the functional connectivity patterns revealed a similar distribution of regions that were also larger on the left than right (data not shown). These results support that the LANG network is distributed across both hemispheres but contains larger regions in the left hemisphere.

### The Candidate Language Network is Similarly Organized and Closely Juxtaposed with Other Association Networks

In all subjects, the LANG network contained regions distributed in multiple zones of association cortex with a broad organizational pattern that paralleled other distributed association networks (Fig. 4). Moreover, the spatial sequence of networks, from LANG to DN-B to DN-A (yellow-pink-red networks in Fig. 4), can be observed in multiple distributed zones in each individual. Clear examples can be seen in temporal and parietal cortices but also along posteromedial cortices, where the LANG network contains a small region in the dPMC neighboring the large regions characteristic of the default network (see S1, S2, S4, S5, and S7 in Fig. 4). Within the IFG, regions of DNB and LANG networks were closely interdigitated, occupying alternating regions curving along the inferior edge of the left IFG in a caudal to rostral axis (see S1, S2, S3 and S5 in Fig. 4 for clear examples). Along the pSTC, DN-B and LANG regions were also closely situated with complex demarcations, in some cases along the length of the superior temporal sulcus (S1, S2, S5, S7 in Fig. 4). In some cases, DN-A regions also bordered LANG regions, for instance near the left IFG (see S2, S5 in Fig. 4), the left TP (S2, S3, S5), the left pSTC (S5, S6), and left dPMC (S5, S7).

The LANG network also bordered the frontoparietal control networks in multiple (but not all) locations. In the IFG, several subjects displayed close-knit LANG and frontoparietal control network regions, particularly FPN-B (see S1, S5, S6, S7 in Fig. 4). LANG and FPN regions were closely positioned along the midline near the pSFG, which also contains a characteristic frontoparietal control network region (e.g., see Fig. 2 in Vincent et al. 2008). The LANG network also bordered the salience (SAL) network near the anterior inferior parietal lobe close to the sylvian fissure and supramarginal gyrus, as well as in posterior regions of the IFG near or in BA6. However, the parietal FPN-A and FPN-B regions did not consistently border the LANG network at or near the pSTC region.

The overall picture was that language regions were distinct but positioned near to separable association networks, with consistent neighboring relationships across individuals that were evident in multiple cortical locations.

### The Candidate Language Network Responds to Language Task Demands

Figure 5 shows the boundaries of the LANG network in each individual, defined by the unbiased data-driven parcellations, overlaid onto regions showing task activation during a language task contrast collected from the same individuals. The spatial similarity can be clearly observed between the two maps, one defined by functional connectivity and one by regional increases in activity during reading sentences compared to lists of non-words. For each subject a threshold was selected by eye, to allow the topography of regions showing strong and weak task effects to be observed, respecting that data quality is not equivalent in all subjects. No masking of the task activation maps was applied that might accentuate their similarity with the intrinsic connectivity maps.

The resulting maps revealed three key findings (Fig. 5). First, in most subjects the regions showing strong task effects were largely confined to the boundaries of the intrinsically defined candidate language network (but see descriptions of exceptions below).

Second, in many places the regions showing task activation had boundaries that occurred at the boundary of the intrinsic LANG network regions. As a particularly striking example, note that in S5 the task activated regions, particularly in the IFG and lateral temporal cortex, almost entirely fill in the spaces between the boundaries of the LANG network. Other clear examples include the left pSTC regions in S1 and left lateral frontal and temporal regions in S6. Third, the association between task activation and intrinsic connectivity was typically not restricted to one part of the brain. Instead, evidence of task activation was found in intrinsic network regions distributed across all 9 zones highlighted in Fig. 1, particularly when all subjects are considered together. For examples, note the small dPMC region of the LANG network in S2, S4 and S7, or the multiple regions on the right hemisphere lateral surface in S2, S6 and S7. The importance of this is that it suggests that the whole distributed network is recruited during the language task contrast, even smaller regions predicted by functional connectivity, rather than just the classical perisylvian language regions.

The overlap was not perfect. Regions of clear task activation that did not overlap with the intrinsic network could be observed in some subjects. For example, in addition to the LANG network regions, the task activation map for S5 (Fig. 5) revealed midline regions along the retrosplenial and posterior cingulate cortices, the anterior medial prefrontal cortex and a circumscribed region of the medial temporal lobe, in an organization reminiscent of DN-A (Braga and Buckner 2017; Braga et al. 2019). Similarly, in addition to the regions of the LANG network, S4 showed regions at or near the primary visual cortex, intraparietal sulcus and frontal eye fields, that typically form part of the dorsal attention network (Corbetta and Shulman 2002). Importantly, the evidence for the recruitment of these other systems was restricted to one or few subjects, while the evidence for a close association between language task activation and the intrinsic candidate language network was evident in all subjects. One possible exception was the left angular gyrus, which showed strong task-driven activation in multiple subjects (e.g., S1, S2, S5; Fig. 5) but did not seem to contain a region of the LANG network defined by intrinsic connectivity in any subject.

To quantitatively test for the selectivity of task-driven responses, the average language task activation effect (mean beta value) was calculated within each of the 6 networks (LANG, DN-A, DN-B FPN-A, FPN-B and SAL) defined *a priori* using functional connectivity (Fig. 6). In all subjects, the intrinsic LANG network showed the highest level of activation during the language task. In most subjects, the LANG network showed a striking degree of selectivity, being activated considerably more than all other networks. In some subjects (e.g., S2, S4 and S7 in Fig. 6) the LANG network was the only network showing activity clearly above baseline. These observations suggest that the LANG network is selectively recruited during the present task contrast involving semantic and syntactic processing. In contrast, neighboring networks showed limited if any evidence of activation, despite their close spatial proximity in multiple cortical zones. One exception was DN-B in S1, which also showed a strong task-activation effect, however this observation did not generalize to other subjects. In S5, DN-A also showed evidence of response that was not found across subjects.

### The Language Network Abuts an Intermediate Network that is Adjacent to Tongue Motor and Auditory Regions

The proximity of Broca’s area to motor representations of the tongue, lips and other oral structures in the inferior portion of the motor strip has been previously noted (Geschwind 1970; Krubitzer 2007). Given the possibility of delineating neighboring functional regions with precision in individuals, we explored the relationship between the language network and sensory and motor regions important for hearing and vocalization. The language network defined in the present set of individuals contained two frontal regions, one in the IFG and one in the pMFG, that were close to the motor strip along the central sulcus (Figs. 1, 4 and 5; see also Glasser et al. 2016; Fedorenko et al. 2010). In addition, a large extended regional response belonging to the LANG network was located in the temporal cortex near to auditory cortex along the supratemporal plane. Seed-based functional connectivity was used to explore the relationship between these LANG regions and the nearby anatomy in the two subjects that provided a motor localizer task (S1 and S2).

We began by mapping orofacial motor and separately auditory regions. Tongue motor regions occupied inferior portions of the central sulcus and nearby gyri (blue regions in Figs. 7A & 8A; see also Carey et al. 2017; Brown et al. 2008; Hesselmann et al. 2004). Functional connectivity from a seed placed in the central sulcus on the contralateral hemisphere revealed a bilateral motor network (MOT; Figs. 7B & 8B). A close correspondence was observed in both subjects between the intrinsic connectivity MOT network and task-driven activations (see also Fig. 6 in Gordon et al. 2017). To define auditory sensory regions, a seed was placed in the contralateral hemisphere on the supratemporal plane at or near Heschl’s gyrus. This defined an auditory network (AUD; Figs. 7C & 8C) based on intrinsic connectivity that comprised a bilateral set of circumscribed regions at the approximate anatomical location of Heschl’s gyrus in both subjects. No auditory localizer was available for these subjects, so the function of the AUD network was presumed based on the bilateral supratemporal distribution of the regions.

**Figure 7:**
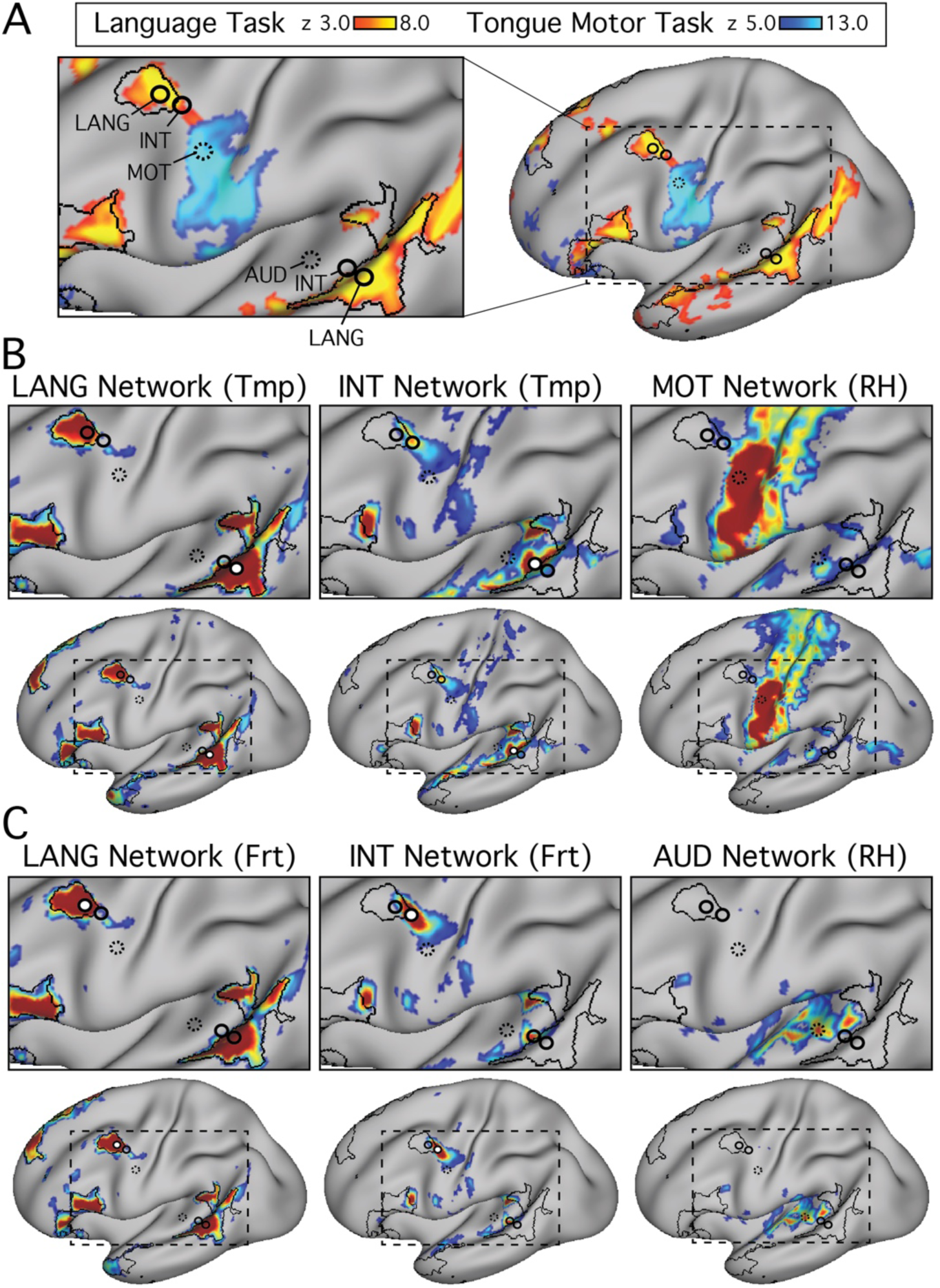
Distributed networks link language regions with tongue motor and auditory regions in S1. An intermediate network (INT) was observed which sits inbetween the language network (LANG) and both the temporal auditory (AUD) and frontal orofacial motor (MOT) regions. (A) Yellow regions show activations during the language localizer task (as in Fig. 5; sentences > non-words), while blue regions show regions displaying increased response during a separate tongue movement task contrast (tongue movements > hand and foot movements) provided by the same subject. The black outline displays the parcellation-defined intrinsic language network (LANG; Fig. 4). Black solid circles are centered on seed vertices that were used to define intrinsic connectivity networks in the remaining panels. The remaining panels show seed-based intrinsic connectivity patterns from seeds selected from the temporal (Tmp; B) and frontal lobes (Frt; C). Auditory and motor regions were recapitulated using functional connectivity using seed regions placed in the contralateral (right; RH) hemisphere as correlation patterns close to the seed are difficult to interpret. Black dashed circles refer to the reflected location of the contralateral seeds. White-filled circles denote the location of the seed used to define correlation patterns in that panel. The INT network displays an organization that parallels the LANG network, containing neighboring regions in both inferior frontal and temporal cortices, as well as along the posterior superior frontal midline (not shown). The function of the INT network is unclear, however its distributed organization and juxtaposition with LANG, MOT and AUD regions in multiple locations suggests it may form part of a hierarchy linking language and sensorimotor functions. Task activations are shown as mean z-normalized beta values, and intrinsic correlations as Fisher’s r-to-z normalized Pearson’s product-moment correlations, ranging from 0.2–0.6, as in Fig. 1.

**Figure 8:**
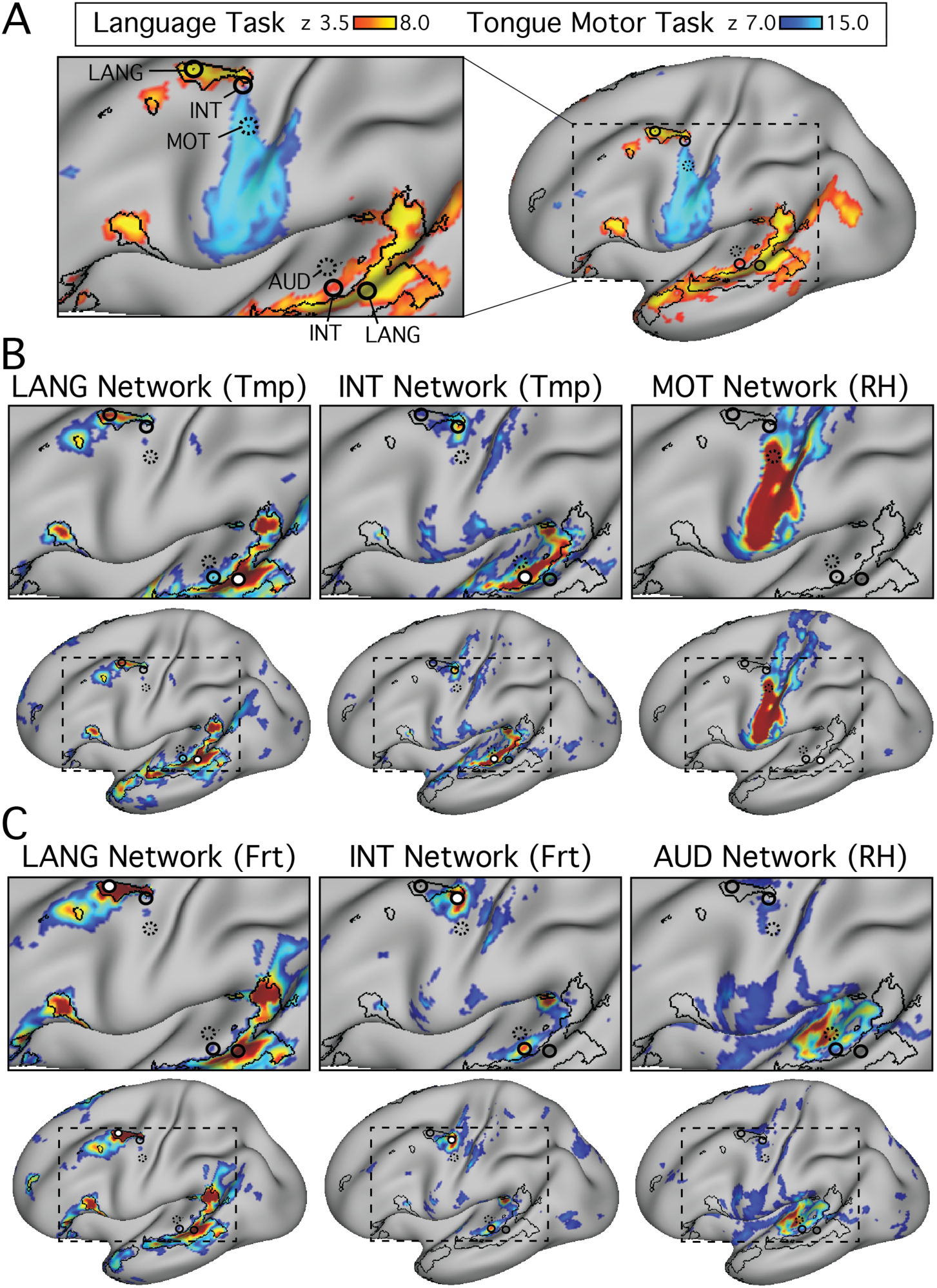
Distributed networks link language regions with tongue motor and auditory regions in S2. Generalizing the findings from S1 (Fig. 7), intrinsic connectivity in S2 also revealed an intermediate (INT) distributed system that bridged the spaces between the language network (LANG) and sensorimotor regions for hearing (AUD) and tongue movements (MOT). A) Task-activated regions are shown for the language (yellow) and tongue motor localizer (blue) task contrasts. The remaining panels show seed-based intrinsic connectivity patterns from seeds selected in the temporal lobe (Tmp; B) and the frontal lobe (Frt; C), as well as in homologous regions of the right hemisphere (RH). Task activations are shown as mean z-normalized beta values, and intrinsic correlations as Fisher’s r-to-z normalized Pearson’s product-moment correlations, ranging from 0.2–0.6, as in Fig. 1.

We next mapped the immediately adjacent zones of the cortex. We hypothesized that the language network regions in the lateral frontal cortex would be juxtaposed with the tongue motor region (Krubitzer 2007) and that the temporal regions would be juxtaposed with the auditory regions. Instead, we unexpectedly observed a small gap between sensorimotor (MOT and AUD) networks and the LANG network near the pSTC and pMFG, and a larger gap in the IFG (Figs. 7 & 8). When seed regions were placed in the spaces between these networks, we identified a smaller, intermediate network (INT). The INT network occupied regions in between the LANG network and the MOT and AUD networks in both frontal and temporal lobes, and also contained a small region neighboring the LANG region in the pSFG. Both subjects displayed a similar distribution of the INT network. Notably, in the frontal lobe the INT network bridged the space between tongue regions and the pMFG LANG region, forming a LANG–INT–MOT sequence of regions. The IFG region did contain a neighboring INT network region (clear in S1 in Fig. 7, less clear in S2 in Fig. 8), however this was separated from the tongue region by the salience network in these subjects (see SAL network in IFG in Fig. 4). Along the midline, a LANG– INT–MOT sequence could also be seen extending from rostral to caudal regions near the pSFG in both subjects (data not shown). In the superior temporal cortex, the sequence of LANG–INT–AUD networks occurred in two separate places, one more caudally near or at the planum temporale, and one more rostrally nearer Heschl’s gyrus (Figs. 7C & 8C).

## Discussion

The present results demonstrate that a distributed language network can be defined within individuals using intrinsic functional connectivity. Organizational details suggest that the network i) is distinct but spatially adjacent to the default and frontoparietal control networks throughout the cortex, ii) has a distributed spatial motif that parallels other association networks, iii) involves upwards of 9 cortical regions in the left hemisphere alone, some of which extend beyond the classical language zones and have not been previously emphasized, and iv) responds in an anatomically-specific manner to language-task demands with adjacent networks showing minimal or no response. We also observed a smaller distinct distributed network that occupies regions in between the language network and the orofacial motor regions in the frontal lobe and the auditory reception regions in the temporal lobe, suggesting a network hierarchy linking language to functionally-related sensorimotor regions. We discuss the implications of these collective observations for understanding the relationship of the language network with the multiple parallel networks that populate association cortex.

### The Language Network Can Be Resolved Within Individuals Using Functional Connectivity

A distributed network that contains regions in classic perisylvian language areas was observed in all 7 individuals tested using intrinsic functional connectivity (Fig. 1; see also Hampson et al. 2002; Lee et al. 2012; Hacker et al. 2013; Glasser et al. 2016). The network was confirmed across analysis methods (Figs. 1, 2 and 4), independent datasets within the same individual (Fig. 3), and could be detected by initiating network definition from multiple distributed locations (Fig. 2). The language network occupied regions that were juxtaposed with other association networks, such as the default, frontoparietal control and salience networks (Fig. 4). The close spatial relationship between neighboring networks, some of which were finely interdigitated (e.g., see sequential LANG and DN-B network regions along the left IFG in Fig. 4), indicates why some prior studies of functional connectivity, especially data-driven methods using group averaged data, may have failed to separate the language network from nearby systems like the default network (e.g., Yeo et al. 2011; Power et al. 2011; but see Mineroff et al. 2018; Blank et al. 2014) and also why studies capturing the network may miss its functional significance.

### The Language Network Parallels the Organizational Motif of Other Association Networks

An intriguing observation of the present study is that the language network is just one of multiple similarly organized distributed association networks. The literature has most often focused on specialization of language regions without consideration of how language networks are similar or dissimilar from other distributed association networks. Our results are fully consistent with a highly specialized left-lateralized network but also illustrate that the distinct language network is just one of several distributed association networks that share a common organizational motif.

Specifically, the network included classical language regions in the frontal and temporal cortices (IFG, pSTC, pSFG, TP, pMFG) as predicted by clinical and task-activation studies (see *Introduction*). However, the language network also extended beyond the classical language system (see Lee et al. 2012; Hacker et al. 2013). Regions were observed in the parietal (dPMC and possibly pSTC region), midcingulate (MCC), and inferior temporal (aITC) cortices, with potentially a further region within the ventromedial prefrontal cortex (Fig. 1 and 4). An anterior prefrontal region (aSFG) was also detected that appeared to be distinct from the pSFG region. Further regions were detected in the right hemisphere, and these regions again displayed a distributed organization that was in many ways homologous to the spatial distribution observed in the left hemisphere (Fig. 4).

When considered together, the resulting language network parallels the distributed motif characteristic of association cortex in the non-human primate (see Fig. 4 in Goldman-Rakic 1988; Margulies et al. 2009; Buckner and Margulies 2019; see also Ghahremani et al. 2017) and previously observed across multiple association networks in humans (Yeo et al. 2011; Power et al. 2011; Margulies et al. 2016; Braga and Buckner 2017). Consistent with earlier observations focused on frontal cortex (Fedorenko et al. 2012), the language network contained side-by-side regions with other well-characterized networks such as the default network, which sits at the apex of a sensory to transmodal cortex hierarchy (Margulies et al. 2016; Buckner and Margulies 2019; Buckner and DiNicola 2019). Neighboring language and DN-B network regions were observed in multiple cortical zones (Fig. 4). The present characterization further illustrates that the spatial juxtapositions are present for multiple distributed components of the language network across the cortex.

For example, the default network contains distributed regions along the posterior, middle and anterior cortical midline, including within the posterior cingulate and retrosplenial cortices, and along the frontal midline (see Zones 5–9 in Fig. 3 in Braga and Buckner 2017; and detailed anatomy in Braga et al. 2019). The language network regions were observed within each of these zones (Fig. 1), with regions reliably defined within the posterior (dPMC; Zone 5 in Braga and Buckner 2017), middle (MCC; Zone 6) and anterior cortical midline at the pSFG (Zone 7), aSFG (Zone 8), and potentially ventromedial prefrontal cortex (Zone 9). Along the lateral surface, language regions were also observed near to or directly bordering the DN-B in the 4 zones highlighted in Fig. 3 of Braga and Buckner 2017, including the IFG, aSFG and pSFG, TP and pSTC. The posterior parietal pSTC region also bordered the prominent default network regions in the inferior parietal lobule (Fig. 4).

The side-by-side relationship between the language network and other distributed association networks could only fully be appreciated when the smaller midline regions were resolved within individuals. This reinforces the notion that the association cortices are organized into parallel, distributed networks, and that in this sense the language network is a characteristic association network.

### Task Activation is Highly Selective for the Language Network

By collecting data during a language localizer task performed by each of our volunteers, we were able to test the hypothesis that the language network, as defined by intrinsic connectivity, is activated by language task demands and also explore the anatomical specificity of the response (see also Glasser et al. 2016). Overlap between connectivity and task activation maps was observed throughout the cortical mantle (Fig. 5). In many cases, the idiosyncratic shape of language network regions closely matched task activated patterns (e.g., see S1, S2, S5 and S6 in Fig. 5), despite being defined in independent data and based on different analysis principles. Importantly, this correspondence extended beyond the classical language regions and often included the smaller regions of the language network. Notable examples include the dPMC region in S2, S4, S5 and S7, the aSFG region in S1, S2, S4, S5, S6, the aITC region in S1, S5, S6, and even the ventromedial prefrontal cortex region in S1, S5, S7 (Fig. 5). The small MCC region showed evidence of task activation in S5 (Fig. 5) at the thresholds selected.

The finding of task activation in these smaller midline regions shows that, under certain task conditions, the entire distributed network is recruited simultaneously in a coordinated and selective manner. In other words, the domain-specialized module appears to be the distributed network, not simply localized regions (see also DiNicola et al. 2019).

The correspondence between functional connectivity and task activation has been noted before (e.g., Smith et al. 2009; Glasser et al. 2016; Gordon et al. 2017; Buckner et al. 2008; Ji et al. 2019). Recently, Tavor et al. (2016) showed that functional connectivity can predict idiosyncratic task activation patterns across individuals. Glasser et al. (2016) also showed that language activation patterns can be recapitulated by intrinsic connectivity using seeds placed in the pMFG and pSTC (see also Hampson et al. 2002). Here we provide corroborative and also additional evidence for spatial specificity. When the average task activation effect was calculated for 6 distributed networks identified *a priori*, the language network showed robust and selective response during the language task (Fig. 6). This was despite that the other networks often possessed regions closely positioned near to the language network simultaneously in multiple cortical zones.

### An Intermediate Network Abuts the Language Network as well as Orofacial Motor and Auditory Regions

Motivated by the hypothesis that the location of prominent language network regions may be explained by their proximity to orofacial motor and auditory regions, we explored the functional anatomy of these regions in two individuals (Figs. 7 and 8). Rather than being juxtaposed, we unexpectedly found a slight separation between sensorimotor regions and the language network in both frontal and temporal cortices. When the functional anatomy of this gap was explored using a seed-based approach, we observed a distinct ‘intermediate’ (INT) network that had a distributed organization and occupied neighboring cortical regions to the LANG network in both lateral frontal and temporal cortices (Figs. 7 and 8), as well as along the dorsal posterior frontal midline. In the frontal lobe, the INT network bordered the LANG network at both dorsolateral (pMFG), dorsomedial (pSFG) and ventrolateral (IFG) locations. The motor task only included tongue movements and it was not possible to map out motor regions involved in the movement of other articulators (lips, pharynx) or the vocal folds (Carey et al. 2017; Brown et al. 2008; Hesselmann et al. 2004; see also Petrides et al. 2005). Hesselmann and colleagues (2004) previously showed that lip movements activate a motor region more dorsal than ventral motor regions activated for tongue movements. One might speculate that the pMFG and IFG INT network regions are associated with different laropharyngeal movements related to independent aspects of articulation and vocalization (see Fig. 3 in de Heer et al. 2017)

The spatial relationships raise the possibility that the LANG and INT networks form a sequence of functional regions that is repeated in multiple cortical zones. The sequence links language regions with tongue movement regions (LANG–INT–MOT) in pMFG (Figs. 7B and 8B) and with auditory regions (LANG–INT–AUD) in the temporal lobe (Figs. 7C and 8C). In posterior IFG, the sequence did not terminate in tongue motor regions, but seemed to lead to the salience network (LANG–INT–SAL; see Fig. 4). The result is a parallel sequence of distributed networks that fall along a gradient from language regions to sensorimotor and possibly other association networks (see also Margulies et al. 2016; Buckner and Margulies 2019; Braga and Buckner 2017; Power et al. 2011).

Following the sequence into transmodal cortex, the LANG network also displayed regions neighboring DN-B in many cortical zones (Fig. 4). In particular, DN-B contains a region in anterior IFG that is closely interdigitated with the LANG network region and extends the sequence into anterior IFG (i.e., DN-B– LANG–INT–MOT). Similarly, a DN-B region is found in the inferior parietal cortex, at or near the temporoparietal junction, which also can be seen as an extension of the sequence into the parietal lobe (i.e., DN-B– LANG–INT–AUD). Altogether, these observations situate the LANG network as falling along a gradient of distributed networks that link auditory and motor cortices with transmodal cortices that support higher-level cognitive functions.

### Conclusions

The present study extends our understanding of the language network by showing that the distributed organization of the language network closely parallels that of other association networks. We reveal the close spatial relationships between language network regions and other distributed systems in classic language regions, and show that the language network sits within a large-scale gradient linking sensorimotor and higher-level association networks. We also resolve small language regions in both hemispheres that have not been previously emphasized and show that these are also language-responsive. The close correspondence of the language network defined by functional connectivity and task-activation suggests that precision functional mapping could aid applied endeavors targeting the language network such as intracranial neuromodulation or to limit complications from surgical resection. Such an approach might be particularly useful for clinical populations that may be unable to perform tasks in the scanner.

## Acknowledgements

We thank the Harvard Center for Brain Science neuroimaging core and FAS Division of Research Computing for support. A. Youssoufian, H. Becker, E. Phlegar and M.K. Drews assisted in data acquisition. R.M. Hutchison assisted with task coding and data acquisition. A. Rodman kindly provided illustrations that were used as cues in the motor localizer task. H. Hoke, T. O’Keefe, R. Mair and S. McMains assisted with data processing optimization. The multi-band EPI sequence was generously provided by the Center for Magnetic Resonance Research (CMRR) at the University of Minnesota. We thank E. Fedorenko for stimuli and thoughtful discussion.

## Grants

R.M.B. was supported by Wellcome Trust grant 103980/Z/14/Z and NIH Pathway to Independence Award grant K99MH117226. L.M.D. was supported by National Science Foundation grant DGE-1745303 (opinions, findings and conclusions expressed in this material are those of the authors and do not necessarily reflect the views of the National Science Foundation). This work was also supported by Kent and Liz Dauten, NIH grant P50MH106435, and Shared Instrumentation Grant S10OD020039.

## Disclosures

The authors declare no competing financial interests.

